# Impact of variability in cell generation times on cell-to-cell variability of protein concentrations

**DOI:** 10.64898/2026.04.23.720286

**Authors:** Syed Yunus Ali, Ashok Prasad, Abhyudai Singh, Dibyendu Das

## Abstract

The influence of arbitrary randomness in cell division times on the variability of protein copy numbers within a lineage ensemble has been recently studied, going beyond the contributions of noisy gene expression and partitioning error. However, variability of protein *concentrations* need separate study, since cell size growth between cell divisions dilute protein concentrations at the same rate as size growth, which also determines mean division times. Here for a model of bursty protein production, we present exact moments (of all orders) of protein concentrations in the cyclo-stationary state, comparing: (i) population and lineage cell ensembles, and (ii) statistics at different cell ages. Two interesting results emerge. While the variance of protein concentration changes with the degree of division time heterogeneity at any cell age, the age-averaged variance is independent of it within lineage ensemble but stays dependent within population ensemble. The skewness within population ensemble is higher in younger cells than within lineage ensemble, and this behavior reverses at older ages. Such a feature vanishes for the age-averaged distribution, with population based skewness always dominating over that of lineage. We also show that mother-daughter correlations in generation times, do not add any significant difference to the results.

## I. INTRODUCTION

Cell-to-cell variability in protein quantity often play vital role in biomedical problems. For example, a drug therapy aimed at stopping irregular cancerous growth may succeed for most target cells, yet may fail eventually and lead to drug resistance due to high expression levels of certain resistance markers in a very small percentage of target cells [1–3]. Such stochastic expression also contributes to microbial persistence [4, 5] and viral replication [6]. Quantitative measure of the degree of gene product variation in cell populations has been possible through technologies of single-cell RNA sequencing and various fluorescent imaging methods to monitor mRNA and protein levels [7–13]. Alongside, understanding the complex regulatory networks [14–16] that influence the variability in the single-cell transcriptomic and proteomic data generated through measurements is necessary for therapeutic benefit.

The regulatory process of a gene may include the altering of its translationally active state(s) by nucleosome and transcription factor binding-unbinding to its promoter, autoregulatory feedback by synthesized proteins, RNA splicing, and post-transcritional regulation by micro-RNAs [17–25]. These coupled stochastic chemical processes which contribute to the overall noise in transcription of mRNA and translation of protein is ‘intrinsic’ to every gene. A lot of theoretical model building has happened to understand analytically, stochastic gene expression within a cell cycle [19, 26–33].

However if the variability of quantity of gene products is studied in a population of cells after many generations, the global or ‘extrinsic’ noise sources associated with successive cell cycles would significantly contribute too, in addition to intrinsic noise. Errors in partitioning of copy numbers during cell division have been studied [34–37]. Noise is also incorporated through variability in cell cycle times of every generation [38–46]. Furthermore cell size growth and size homeostasis influences and gets influenced by gene expression [38–42, 47– 53]. Cell division itself has been viewed to be controlled by timer, or threshold cell size, or added cell size [54–62]. Thus cell generation times are influenced by cell size control. Experiments are done either with single cell lineages in multiple microfluidic channels, or with exponentially growing cell populations. The statistics are distinct, depending on whether the sampled cells belong to lineage or population ensemble [63–65], and also on the specific cell ‘age’ [46, 66, 67]. Given these multiple layers of complexities, systematic theoretical calculations on standard simple models of protein production disentangling contributions of separate noise sources, is generally useful in enriching our understanding of the underlying processes.

Recognizing the importance of noise induced through cell division, various works looked at cumulants and distributions of mRNA and protein copy numbers in the cyclo-stationary state (after many generations), albeit ignoring the randomness in cell division times completely [33, 34, 63, 68–71]. Yet as noted above, experimental data indeed shows generation time heterogeneity. For example, Erlang distribution was fitted to division times of mouse cells [43], beta-exponential in case of Caulobacter crescentus [59], log-normal in the case of Lymphocyte B-cells [38], and skew-normal in mouse intestinal organoid cells [72]. Another line of work assumed the cell cycle to have *N* stages, each governed by a Poisson process, and thus the division time to have an Erlang or Hypoexponential distribution [53, 63, 73– 75] – various results associated with the ‘age-averaged’ distributions of the copy number of gene products were obtained. However, the problem of general random cell generation times remained theoretically open.

Recently, a framework has been developed to calculate analytically exact cyclo-stationary distributions of copy numbers within lineage ensemble as series forms, for any arbitrarily random (but uncorrelated) cell division times [46]. The separate contributions of the three sources of noise (gene expression, random partitioning error, and division time heterogeneity) to the copy number variability could be segregated. Cell age-specific statistics (at birth, before division, and ages in between), as well as age-averaged ones could be derived. Yet several important experimentally relevant issues remained untreated. What happens to the method in the case of population ensemble? Moreover, cell sizes grow through every cycle and roughly half at division. On one hand the mechanisms to attain homeostatic size distributions introduce mother-daughter generation time correlations [57, 76, 77]. On the other hand, cell size growth makes transcription rates volume dependent, and concentrations dilute even when proteins do not degrade [53]. Although these mechanisms are very hard to treat analytically exactly with simultaneous stochastic variation of cell volume (*V* (*t*)) and copy number (*n*(*t*)), yet is an exact treatment possible for effective description in terms of protein *concentrations* (*x* = *n/V* ), similar to earlier works on gene expression [53, 78–80]?

In fact, a recent work [65] addressed the above issue of cell growth and gene expression coupling by studying the statistics of ‘age-independent’ protein concentrations. They found the interesting results that fluctuations are greater in population ensemble than within lineage, and for lineage the variance is independent of division time heterogeneity. Yet the problem is not completely solved without finding the infinite set of moments, which is equivalent to finding the full distribution in series form like in the above mentioned work [46]. Moreover, without getting into age-specific statistics, various relationships of the population versus lineage ensemble were not realized.

In this paper, we extend the general formalism of [46] for independent random cell division times, to obtain the statistics of protein concentrations in growing and dividing cells. Like [65], we couple the concentration dilution rate to cell growth rate and mean cell division time (see Section II). For both lineage and population ensembles, starting from self-consistent integral equations of distributions, we present exact recursive relations between infinite set of moments, thus formally closing the problem. We do this for new-born cell, cells about to divide, cells at any intermediate age, and also for age-averaged distributions (see Section III). While the results of [65] are reproduced, we have many more results which clarify the subtle relations of cell ages and cell ensembles. In Section III F, we explore how the mother-daughter correlations in division times due to ‘adder’ mechanism of size control, cause very little departure from our analytical results obtained by ignoring correlations. Finally we summarize our results in Section IV.

## II. MODEL COMPONENTS: BURSTY PROTEIN PRODUCTION, DILUTION DUE TO CELL GROWTH, AND RANDOM PARTITIONING AFTER RANDOM CELL DIVISION TIMES

We consider a stochastic model of the evolution of protein concentration *x* = *n/V* in a growing cell [53, 78–80], where *n* is the instantaneous copy number and *V* is the cell volume. As experimentally observed across different cell types, the overall synthesis rate of the given gene product (i.e. number of molecules produced per unit time) scales with cell size, and this scaling ensures concentration homeostasis [81–86]. A schematic of *V* (*t*), *n*(*t*) and *x*(*t*) as function time *t* is shown in Fig. 1. Note that, as the figure indicates, while average ⟨*V* ⟩ and ⟨*n*⟩ is halved at every division, ⟨*x*⟩ in a newborn cell remains same as its mother. We do not study independent stochastic updates of *V* and *n*, but instead a direct stochastic evolution of *x*(*t*) which effectively takes care of protein translation and cell size effects. Further details of the model are provided below.

**FIG. 1.**
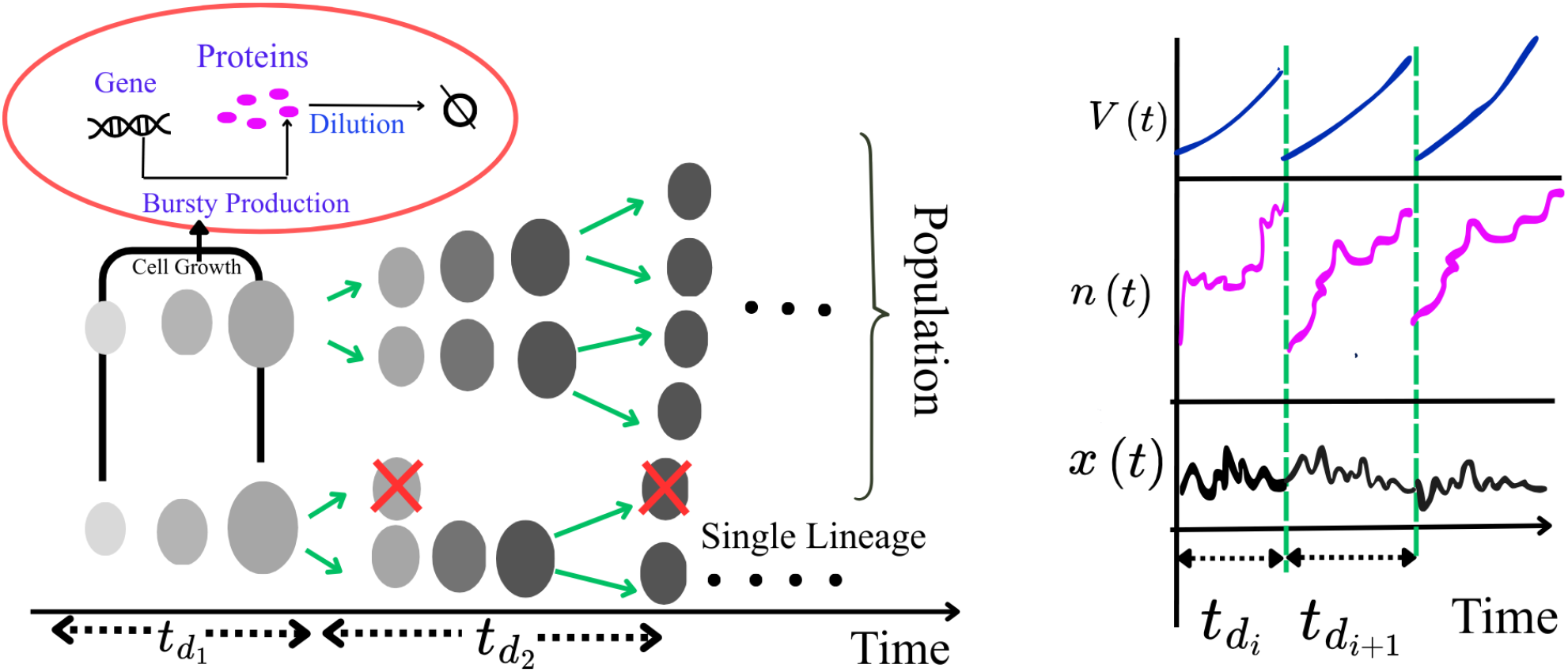
Schematic of cell size growth with rise of protein concentration due to gene expression within a cell cycle, depicted by gradual darkening of grey shade. The concentrations don’t change on an average at the time of cell division (see the colors). In the lineage ensemble (bottom) one of the two lines of cells are discarded after every division. In the population ensemble (above) cell lines of all generations are retained. The random generation times are denoted by 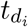 and for *i* ≫ 1, cyclo-stationarity is attained. On the right, a schematic sketch of cell volume *V* (*t*), copy number *n*(*t*) and concentration *x*(*t*) of proteins, is shown as a function of time – big jumps of *x*(*t*) are absent at cell divisions, unlike *V* (*t*) and *n*(*t*).

- Proteins are produced in bursts, under the assumption of very fast relative degradation of mRNA compared to proteins [21, 29, 78] – as a result, protein concentration augments as (*x* → *x* + Δ*x*) with an exponentially distributed (i.e. 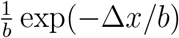) burst size Δ*x*, having mean ⟨Δ*x*⟩ = *b*. The variation of concentration within a cell cycle is indicated by a color gradient for the first two cycles in Fig. 1, but it stabilizes around a mean soon as shown in the *x*(*t*) vs *t* plot alongside. The transcription rate *k* of mRNA concentrations is volume-independent. The forward master equation for the probability distribution of protein concentration *P* (*x, t*) is [78]:

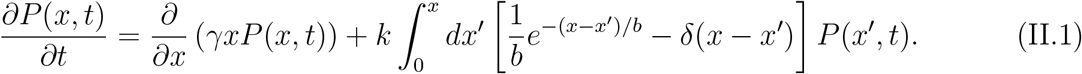 In the above, the proteins are assumed not to degrade within the relevant cell-cycle times, and yet their concentration decays at a rate *γ* due to an increase in cell volume within a cell-cycle. Consequently, *γ* is related to the mean cell division time as we discuss below.
- As we focus on random cell division times *t*_*d*_ (see Fig. 1), we assume those to have a distribution *g*(*t*_*d*_) and uncorrelated over generations. The protein dilution rate *γ* mentioned above is assumed to be equal to the cell volume doubling rate, and thus *γ* = ln 2*/*⟨*t*_*d*_⟩ where ⟨*t*_*d*_⟩ is the mean division time. To respect this relationship, certain parameters of the distribution *g*(*t*_*d*_) get related to *γ*. For example, the Erlang distribution

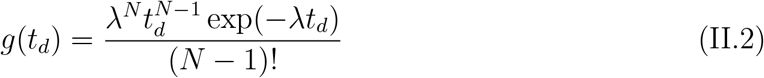

(with *N* = 1, 2, 3…), has a mean ⟨*t*_*d*_⟩ = *N/λ*, and hence *γ* is related to *λ* = *γN/* ln 2. Note that *N* = 1 is an exponential distribution while *N* → ∞ gives a Dirac-delta function – see Fig. 7(a) in Appendix. A. Similarly, for the Beta-Exponential distribution

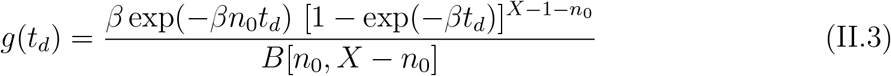

with threshold *X* and growth rate *β*, to have the mean cell-cycle time 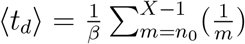 Hence *γ* is related to 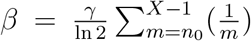. Note that the width of *g*(*t*_*d*_) decreases with increasing threshold *X* (see Fig. 7(b) in Appendix. A). Although we present general results for any *g*(*t*_*d*_) in this paper, we study below Eqs. II.2 and II.3 as specific examples.
- As we are interested in the statistics of protein concentrations after multiple cycles of cell division, we need to discuss next how to partition the concentration at any cell division. We note that after division, both number *n* and the volume *V* are shared, on average, equally by two daughters. Hence mean concentration does not change during division (see Fig. 1). But due to the binomial partitioning error of the copy number, and slight volume inequality to the daughter cells, it is expected that partitioning noise will add to the protein concentrations. Thus, we assume *x*_−_ (concentration just before division) → *x*_+_ = *x*_−_ ± *δx*, where *x*_+_ is the concentration of proteins just after birth. We assume that the partitioning noise *δx* is ‘normally distributed’ with mean = 0 and variance *σ*^2^ = *εx*_−_*/*4; the limit *ϵ* → 0 recovers the standard model of bursty gene expression without cell division [78], consistent with experiments [20, 21, 87]. The parameter *ε* (with unit of concentration), a tunable controller of the width of the Gaussian distribution, needs to be small such that *σ* ≪ *x*_−_. Thus although *δx* ∈ (−∞, ∞), in practice it remains quite small and in no case *x*_+_ ever becomes negative. To avoid in principle any negative *x*_+_, the distributions with finite support like box or beta may be chosen. But exact calculations become hard to carry through with them, unlike with a Gaussian as we show below. We perform simulations to show that the results for a box distributed partition error in concentration (with probability 1*/*2*W* over *δx* ∈ [−*W, W* ], and zero otherwise), follow similar trends to that of Gaussian distribution with comparable variance (i.e. with 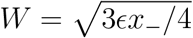), as one would expect.
- The above model is studied for two ensembles in this paper. The first one is *single lineage*, in which one imagines a single copy of cells retained after every division, in several independent microfluidic channels. The protein content of cells in different channels are compared to obtain the statistics. Such ensemble average across multiple channels, is quantitatively equivalent to doing long time average in the steady-state of a lineage within a single channel. The second one is *population* ensemble in which all cells of an exponentially growing population are used to obtain the statistics. Distinct formal equations and results for these two cases will be discussed in the paper. We would like to note that the cell age distributions *ψ*(*τ* ) and *ϕ*(*τ* ) within the lineage and population ensembles respectively, are well known in the literature [66, 67, 88] for arbitrary cell division time distributions *g*(*t*_*d*_):

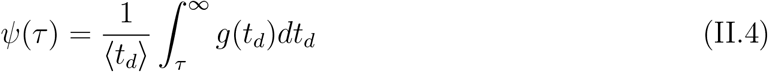

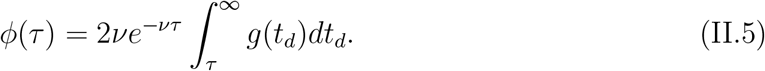 Here, the growth rate *ν* of the exponentially growing population is implicitly given by

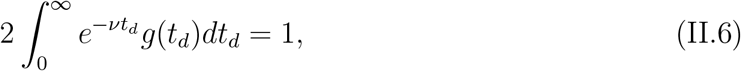

for binary division.
- Although our analytical results are obtained for independent cell generation times, we compare our results with a simulated ‘adder’ model of cell szie control, which is known to have mother-daughter generation time correlation. See section III F for the model and results.

For the model outlined above, we now derive the exact moment identities at various ages, as well as the age-averaged moments, in the cyclo-stationary state achieved after many cycles of cell division.

## III. RESULTS

### A. Statistics of protein concentrations at birth (age *τ* = 0)

The protein concentration *x*_+,*i*_ in a cell at birth in the *i*-th cycle evolves through gene expression and dilution due to the cell growth and reaches a terminal concentration *x*_−,*i*_ just before its division in a random generation time *t*_*d,i*_. Post division the protein concentration is *x*_+,*i*+1_ in one of the daughter cells. On attaining stationarity after several cycles of division (*i* ≫ 1), the distribution of concentration at birth (*x*_+_) stabilizes to 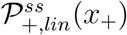 within a *single lineage*, and assuming lack of inter-generational correlation i.e. 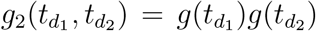, obeys the following self-consistent equation (see Appendix. B):

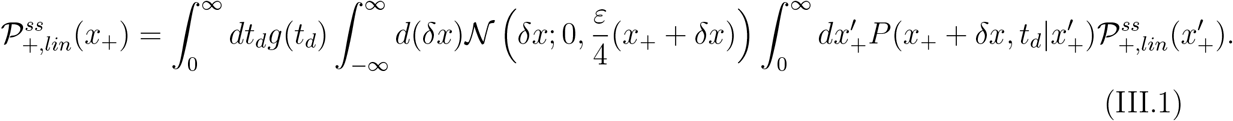

Here, 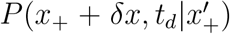 is the probability of the protein concentration being *x*_+_ + *δx* at the end of the cell-cycle starting from 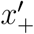 at birth, evolving through the master equation (Eq. II.1) for the bursty production and dilution protein kinetics. N(*δx*; 0, *σ*^2^) denotes the Normal distribution of the partitioning error *δx*, such that the concentration of *x*_+_ + *δx* of the mother produces *x*_+_ of the daughter after division. Note 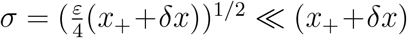 (as discussed above). The Eq. III.1 leads to the equation for the Laplace transform of the concentration 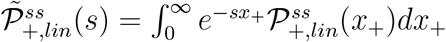 (see Appendix. B):

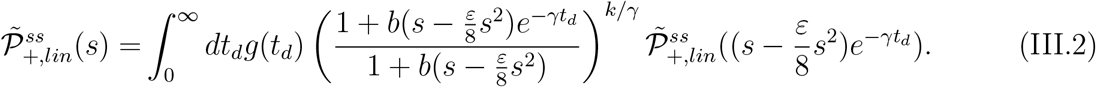

On the other hand, the cyclo-stationary protein concentration in newborn cells within an exponentially growing *population ensemble* has a distribution 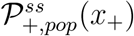 satisfying:

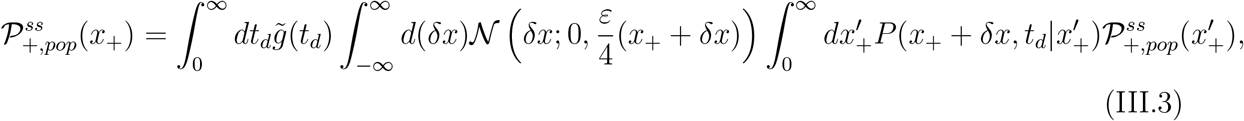

which is similar to Eq. III.1, except *g*(*t*_*d*_) is replaced by

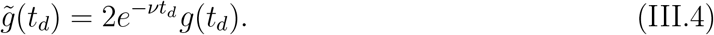

In the above, the factor 2 arises as either of the two daughters may inherit the concentration *x*_+_ from their mother and contribute separately to the population. The distribution 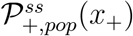 is a ratio *N*_+_(*x*_+_, *t*)*/N* (*t*) of the number of newborn cells (*N*_+_(*x*_+_, *t*)) with protein concentration *x*_+_, and the population size *N* (*t*). The factor 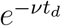 in Eq. III.4 follows from the increase of *N* (*t*) by the factor 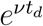 after a cell generation time *t*_*d*_. Recall that *ν* follows from Eq. II.6. Consequently, the Laplace transform of 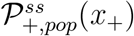 has a similar formula as Eq. III.2 (details in Appendix B) and given by :

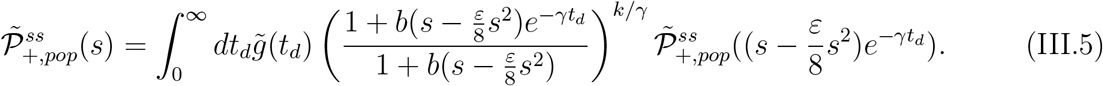

To demonstrate the validity of the Eq. III.2 and Eq. III.5 in Fig. 2, let us take the example of an Erlang distribution (Eq. II.2) of cell division times. In Fig. 2(a) we show the protein concentration distributions for lineage and population ensembles, obtained through Gillespie simulation [89] of the model discussed in the sec. II. The Laplace transform of the latter data have been plotted in symbols in Fig. 2(b). Next, we numerically iteratively solve the self-consistent Eqs. III.2, III.5 and depict those in solid lines in Fig. 2(b) – the match with Gillespie simulation data is clearly seen.

**FIG. 2.**
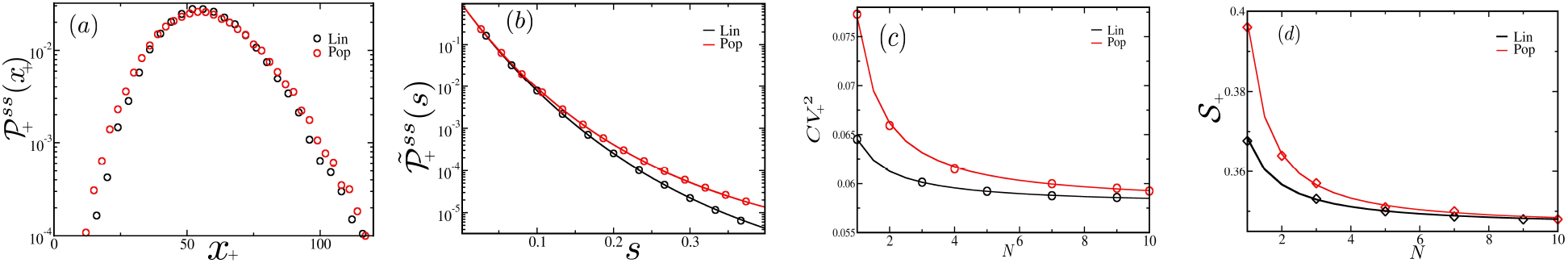
Statistics of protein concentrations (*x*_+_) in new born cells, for population (red) and lineage (black) ensemble respectively. (a) 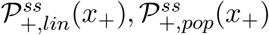 from Gillespie simulation. (b) Laplace transform 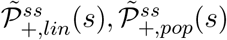 against Laplace variable *s*, solved from Eqs. III.2,III.5. (c) Noise 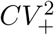 and (d) Skewness S_+_ in gene product levels (*x*_+_) as a function of parameter *N* for the Erlang distribution *g*(*t*_*d*_). In (b-d), solid lines denote analytical results, while symbols correspond to simulation data. The model parameters are *k* = 1 min^−1^, *b* = 2 nM, ⟨*t*_*d*_⟩ = 20 min, *ε* = 4.0.

The problem can be formally analytically solved if the infinite set of all the moments 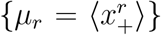 (for the lineage ensemble) and 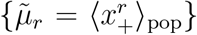 (for the population ensemble) can be obtained, such that

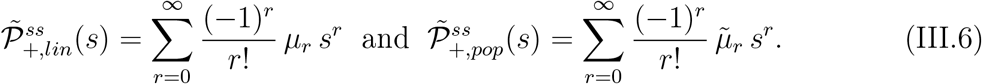

We show in Appendix B that exact recursion relations for moments of any order can be obtained. The final result is given below:

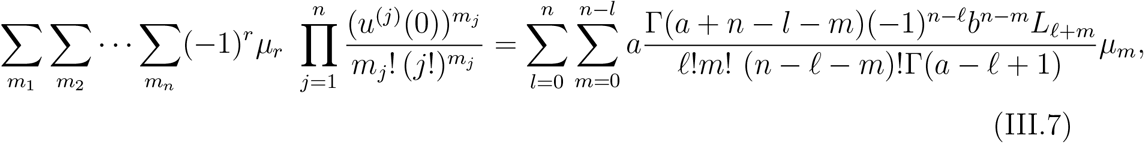

where 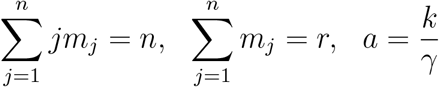, and 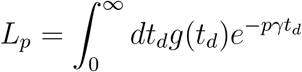,

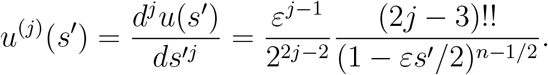

Here {*m*_1_, *m*_2_, · · ·, *m*_*n*_} are non-negative integers satisfying the above two conditions, determining the value of *r* ∈ {1, 2, · · ·, *n*} in different terms. Thus Eq. III.7 gives a linear relation of moments *µ*_*n*_, *µ*_*n*−1_, · · ·, *µ*_1_. A moment of any order can be recursively solved from the system of linear equations. Similarly, a recursion relation corresponding to Eq. III.7 may be derived for the *population* ensemble with 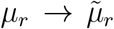 and 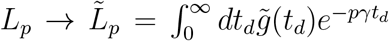, starting from an equation similar to Eq. B5 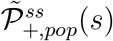. The latter would lead to the for solution of the moments 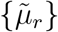.

Using 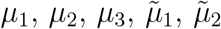, and 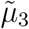 we explicitly provide here the mean 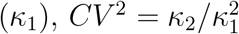, and Skewness 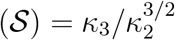 for the lineage and population ensembles, where *κ*_2_ and *κ*_3_ are their variance and third cumulant:

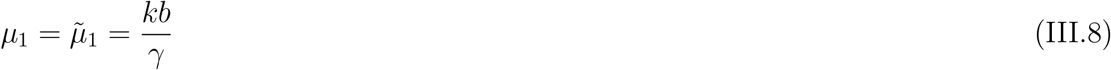

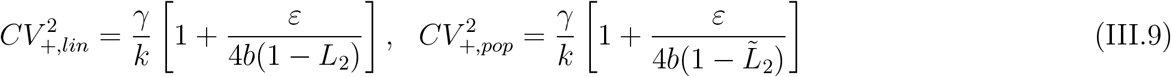

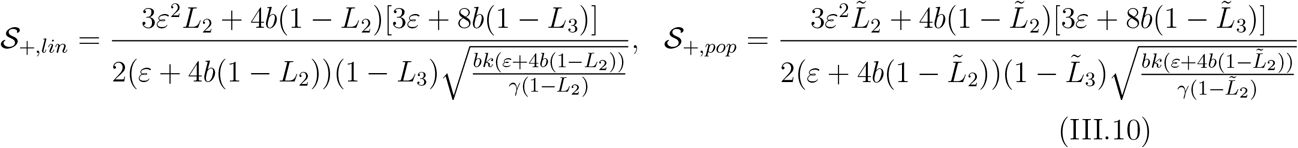

An important thing we observe in Eqs. III.9 and III.10 is that the relative fluctuations depend explicitly on the division time distribution *g*(*t*_*d*_). For the Erlang distribution we can derive the following results. Due to the relationship of the *λ* parameter with the protein dilution rate *γ*, we have 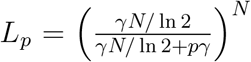, and 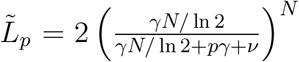 (where *ν* = *λ*(2^1*/N*^ − 1)). By using these expressions in the equations above, we can see that if the parameter *N* of the the Erlang distribution in Eq. II.2 is varied, then the *CV* ^2^ decreases in Fig. 2(c) with increasing *N* (i.e. decreasing width of *g*(*t*_*d*_) as shown in Fig. 7 in Appendix. B). We observe that the coefficient of variation of the population is always greater than that of the single lineage, i.e., 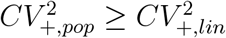 for all *N* (similar as in [65]). This result follows from 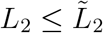. Similarly, in Fig. 2(d) we see that the skewness 𝒮_+,*pop*_ ≥ 𝒮_+,*lin*_, thus the population skewness is always greater than or equal to the lineage skewness.

For the Beta-Exponential distribution given by Eq. II.3, we get 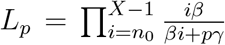 and 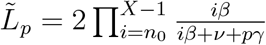 as a function of *X*, which controls the width of this distribution. Here *ν* needs to be obtained numerically using Eq. II.6. Calculating the *CV* ^2^ and the skewness using these expressions, we find again that the population values are greater than or equal to the single lineage values, as shown in Fig. 8(a)-(b) (Appendix. B), where we plot 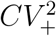 and 𝒮_+_, by varying threshold *X*.

Finally, although we do not have analytical results for partitioning error of concentration *δx* for bounded distributions, we study the case of the box distribution through simulations and present our data in Fig. 9. The 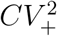 and 𝒮_+_ versus *N* curves in Fig. 9(a)-(b) (Appendix. B), are almost identical to Fig. 2(c)-(d) for the Gaussian error, assuring that our analytical results are well applicable even if the partitioning distribution are non-Gaussian.

### B. Statistics of protein concentrations in cells just before division

The terminal protein concentration is *x*_−_ before cell division and its stationary state distribution is 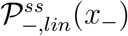. As starting from 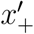 the concentration evolves to *x*_−_ propagated By 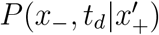, for a *lineage* we have

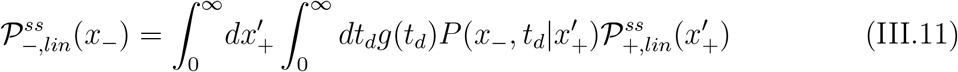

On the hand within a *population*, as 2 cell lines may contribute to *x*_−_ and the cell population *N* (*t*) grows by a factor 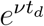 in cycle time *t*_*d*_, the factor 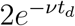 would multiply *g*(*t*_*d*_) (see Eq. III.4), and the concentration distribution would satisfy:

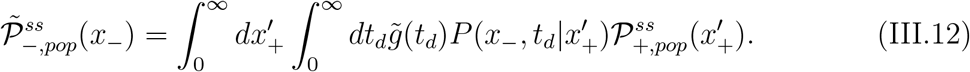

In Fig. 3(a) we plot the concentration distributions 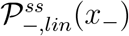 and 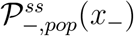 for the two ensembles, obtained from Gillespie simulation. As we know the Eqs. III.2 and III.5 for the Laplace transforms 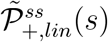 and 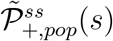, we may show (Appendix C) that the Laplace transforms 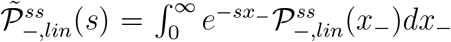 and 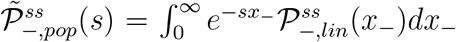 satisfy:

**FIG. 3.**
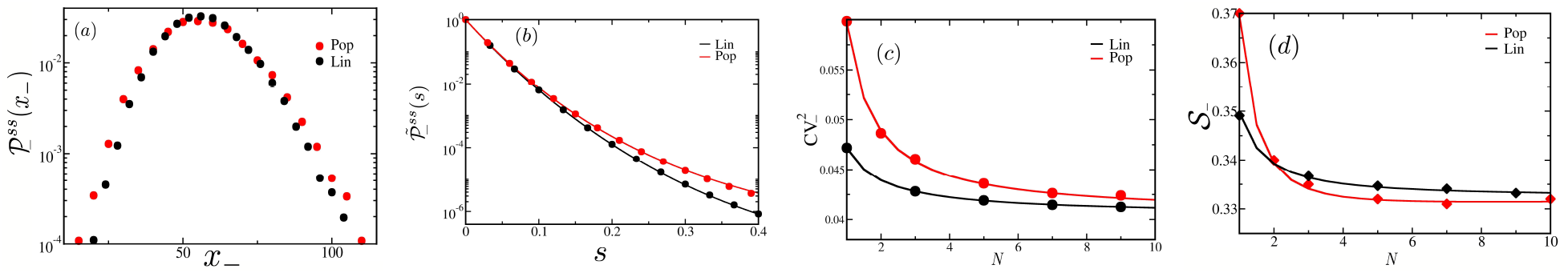
Statistics of protein concentrations (*x*_−_) in cells about to divide, for population (red) and lineage (black) ensemble respectively. (a) 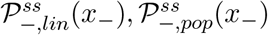 from Gillespie simulation. (b) Laplace transform 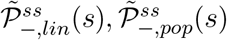 against Laplace variable *s*, solved from Eqs. III.13. (c) Noise 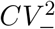 and (d) Skewness 𝒮_−_ in gene product levels (*x*_−_) as a function of parameter *N* for the Erlang distribution *g*(*t*_*d*_). In (b-d), solid lines denote analytical results, while symbols correspond to simulation data. The model parameters are *k* = 1 min^−1^, *b* = 2 nM, ⟨*t*_*d*_⟩ = 20 min, *ε* = 4.0.

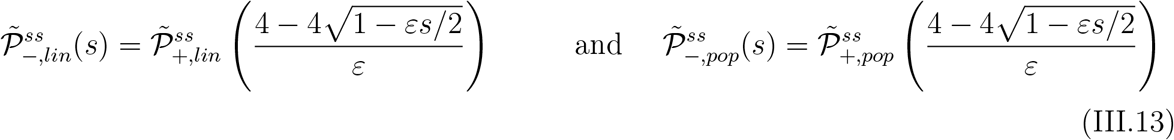

In Fig. 3(b) we confirm the validity of the predicted expressions of 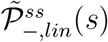 and 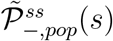 (in Eqs. III.13) based on numerical solutions of 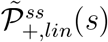 and 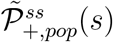 from Eqs. III.2 and III. 5, against the Laplace transforms of the Gillespie data from Fig. 3(a). Furthermore, since we know the series expansion of 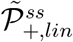 in terms of its moments (Eq. III.7), we immediately obtain the equation for the *n*th moment 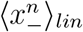 of 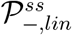 in terms of the moments {*µ*_*r*_} of 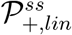:

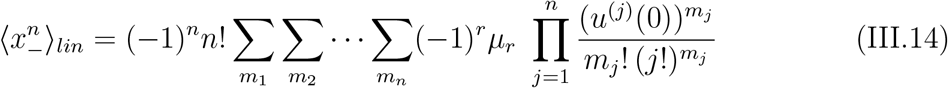

The symbols in the above equation satisfy the same conditions as in Eq. III.7. The *n*th moment of 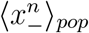 within population ensemble satisfies a similar equation as above with *µ*_*r*_ replaced by 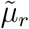. Explicitly, we have the mean, *CV* ^2^, and the skewness 𝒮 in both lineage and population ensembles as follows:

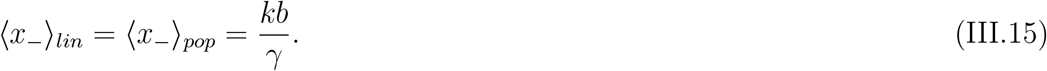

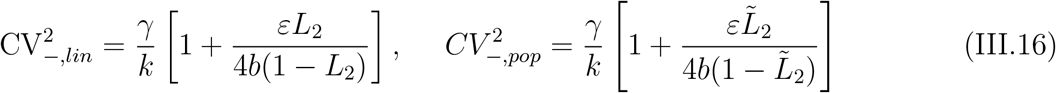

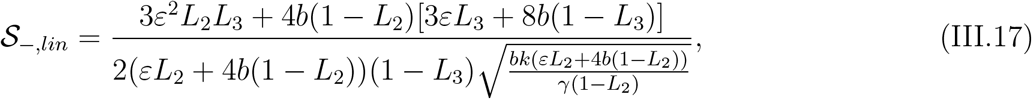

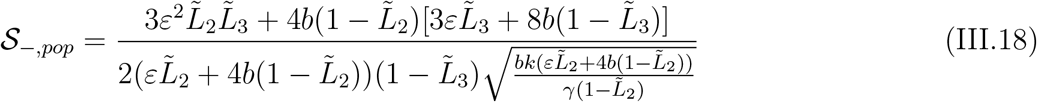

In Fig. 3(c), for the Erlang distributed *g*(*t*_*d*_), we see that 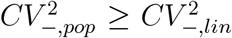 for all *N*, but in Fig. 3(d) we see that there is a crossing of the curves of 𝒮_−,*pop*_ and 𝒮_−,*lin*_. We also demonstrate in Fig. 8 (c) and (d) similar trends of 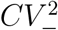 and skewness *S*_−_, for the Beta-exponentially distributed *g*(*t*_*d*_). We will see below that the interesting feature of crossing of skewness curves occur for protein distributions at other non-zero cell ages. We also study the case of the box distributed partitioning error *δx* through simulations – the 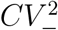 and 𝒮_−_ versus *N* curves in Fig. 9(c)-(d) are almost identical to Fig. 3(c)-(d), confirming the usefulness of our analytical results for Gaussian partitioning error.

### C. Statistics of protein concentrations at any cell age *τ >* 0 before division

The distribution of protein concentrations for cells at a specific age *τ* after birth and before division, is given in terms of the distributions at birth as follows:

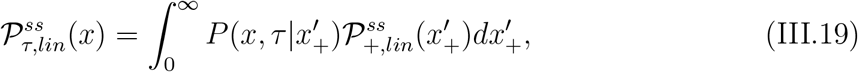

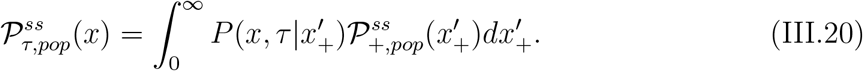

The Laplace transforms of Eq. III.19 and III.20 on substituting 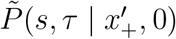 from Eq. (B3), give Eqs. D1 and D2. We then expand the Laplace transforms as a Taylor series in *s* and equating the coefficients, obtain the moments 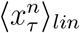 and 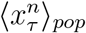 (of any order *n*) of the age-specific distributions in both the ensembles – see Eqs. D3 and D4 in Appendix D. From those general expressions, we specifically have the mean, *CV* ^2^, and skewness 𝒮 for lineage and population ensembles as follows

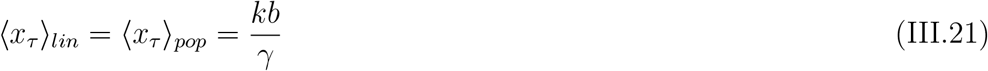

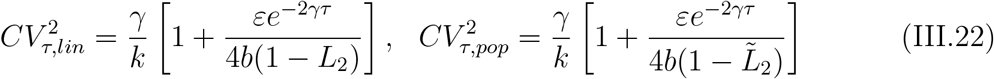

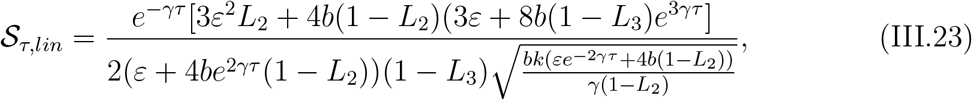

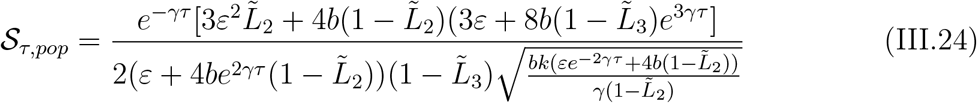

In Fig. 4 (a), we show that for Erlang distributed divison times, just like *x*_*±*_, in the case of *x*_*τ*_ (for any age before division) we have 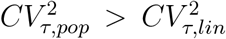. On the other hand 𝒮_*τ,pop*_ crosses from a higher to a lower value with respect to 𝒮_*τ,lin*_ as a function of *N* (of Erlang distribution), for a range of cell ages *τ* – the crossing is seen at *N* ≥ 1, over *τ* ∈ [4, 18) – see Fig. 4(b) for *τ* = 10 for example. For *τ <* 4 we have 𝒮_*τ,pop*_ *>* 𝒮_*τ,lin*_, while for *τ >* 18 we have 𝒮_*τ,lin*_ *>* 𝒮_*τ,pop*_ – this is shown in Fig. 10 in Appendix. D. Varying *N* (that is varying the distribution *g*(*t*_*d*_)) is experimentally difficult. Instead, for a fixed distribution *g*(*t*_*d*_) (with fixed *N* ), it is easier to study the statistics of protein concentrations *x*_*τ*_ of different cell ages. With that aim, we show in Fig. 4(c)-(d) the *CV* ^2^ and 𝒮 as a function of *τ* for *N* = 3. Interesting, we see that 𝒮_*τ,pop*_ crosses 𝒮_*τ,lin*_ also as a function of *τ* for a fixed *N* . Thus, we predict that at early cell ages the skewness of protein concentrations will be higher in the population than in the lineage, while as the cell becomes older, the behavior is reversed. It is thus very interesting to ask what will happen to this property if one looks at concentration distribution averaged over all cell ages. We consider this below.

**FIG. 4.**
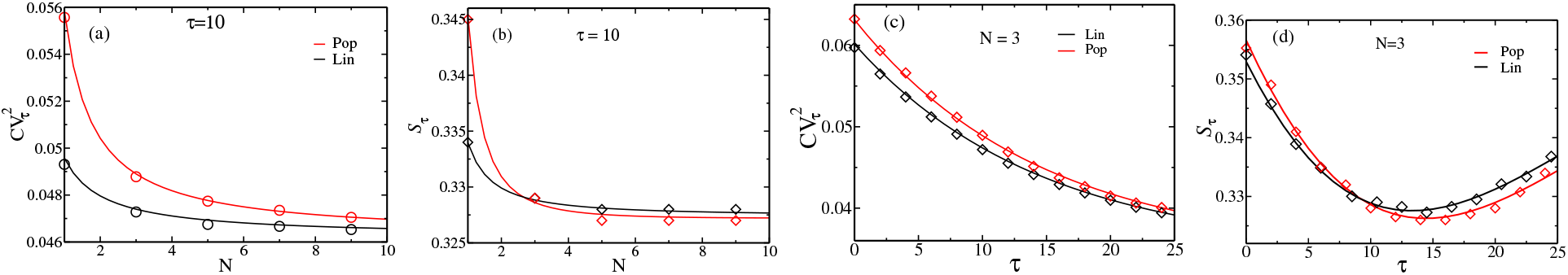
Statistics of protein concentrations (*x*_*τ*_ ) at different cell age *τ* for population (red) and lineage (black) ensemble respectively. Age-specific (a) noise 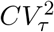 and (b) skewness 𝒮_*τ*_ of gene products as a function of the Erlang parameter *N*, for *τ* = 10 min. (c) Noise 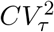 and (d) skewness 𝒮_*τ*_ as a function of varying age *τ*, for *N* = 3. Solid lines denote analytical results, while symbols correspond to simulation data. The model parameters are *k* = 1 min^−1^, *b* = 2 nM, ⟨*t*_*d*_⟩ = 20 min, *ε* = 4.0.

### D. Age-averaged distributions of protein concentrations and moments

To calculate theoretically the distributions averaged over all cell ages, one needs to know the appropriate cell age frequencies *ψ*(*τ* ) and *ϕ*(*τ* ) for any division time distribution *g*(*t*_*d*_), which are given by Eqs. II.4 and II.5. The lineage and population age-averaged protein concentration distribution are thus

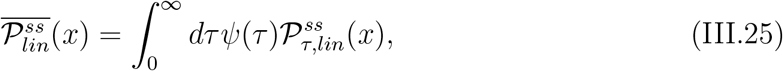

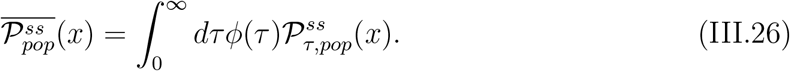

We now use a similar strategy as in the above sections, to calculate the moments. We Laplace transform these equations, then substitute Eqs. D1 and D2, series expand in *s*, and equate the coefficients of *s*^*n*^ from both sides which yield the exact *n*th moments 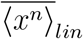 and 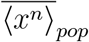 (for any order *n*) of the age-averaged distributions in the two ensembles – see Eqs. E3 and E4 in Appendix E. The cumulants up to third order of of the age-averaged protein concentrations is provided below for both the ensembles:

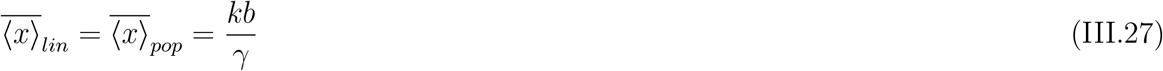

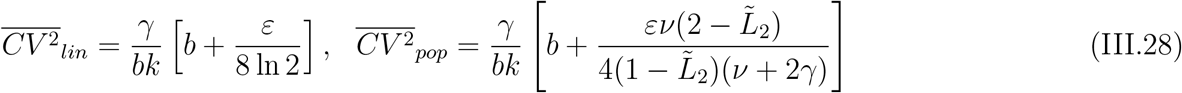

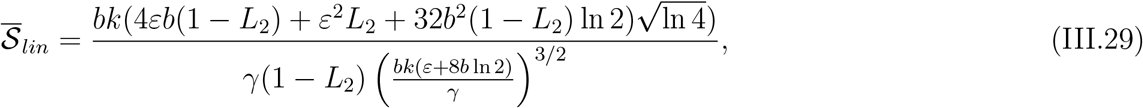

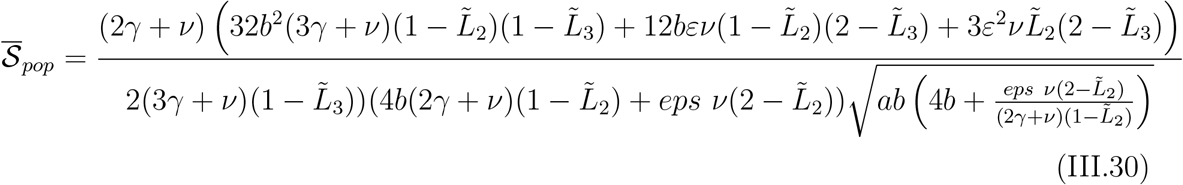

Although 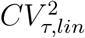 at any cell age *τ* depends on the division time distribution *g*(*t*_*d*_) (see Figs. 2(c), 3(c), 4(a)), in Eq. III.28 we see a fascinating result that the age-averaged 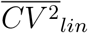 is independent of *g*(*t*_*d*_) (as noted in an earlier publication [65]). In Fig. 5(a) this is demonstrated by comparing with Gillespie simulation data – see the black line for 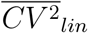 which is constant as *N* varies. In comparison 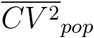 (from Eq. III.28) varies with *g*(*t*_*d*_) through 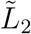– its behavior is shown in Fig. 5(a). Thus the protein fluctuations in the population ensemble remains sensitive to the division time randomness unlike the lineage ensemble, at the age-averaged level too. This is a vital prediction for experiments.

**FIG. 5.**
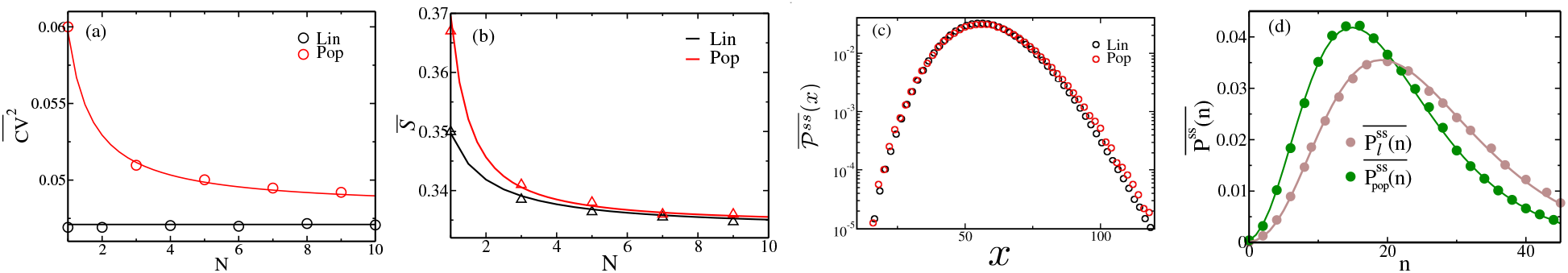
Statistics of protein concentrations 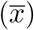 averaged over all cell ages prior to division. (a) Noise 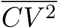, (b) skewness 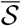 of gene products as a function of the Erlang parameter *N*, and (c) the full distributions 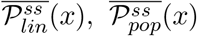 for population (red) and lineage (black) ensemble respectively. (d) The full protein number distributions 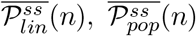 in the two ensembles (in green and brown). Solid lines denote analytical results, while symbols correspond to simulation data. The parameters for the model of concentration are *k* = 1 min^−1^, *b* = 2 nM, ⟨*t*_*d*_⟩ = 20 min, *ε* = 4.0. The parameters for the model of number are *k* = 0.5 min^−1^, *b* = 2, *γ* = 0.1 min^−1^and ⟨*t*_*d*_⟩ = 20 min.

In Fig. 5(b) we show the age-averaged skewness 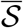 for population and lineage. The fascinating thing is that as a function of *N* there is no crossover of the population and lineage curves as in Fig. 4(b) for an intermediate cell age. Thus it seems that the higher weightage of lower cell ages in age-distributions dominate, and make the age-averaged behavior of 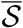 similar as that of young cells (see Fig. 2(d) and 10(a)).

Above, we have obtained formulas of the moments 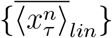 and 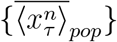 of any order of age-averaged protein concentration distributions 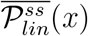 and 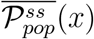 respectively. This implies that we know the exact series expansions of the the Laplace transforms 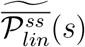 and 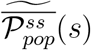. Yet, it is not easy to inverse Laplace transform these series forms and obtain explicitly the distributions 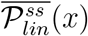 and 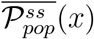 in the concentration domain, although the problem is formally solved through the infinite set of moments. We show numerical plots of the age-averaged conentration distributions in Fig. 5(c) obtained from Gillespie simulations – as the mean concentrations are same in the two ensembles, the differences of these two distributions are only visible if we move considerably away from their centers. Note that the moments are thus more sensitive markers of the difference between the two ensembles compared to the concentration distributions.

In contrast, the full distributions of the protein copy numbers are possible to be analytically calculated explicitly, and they differ significantly for the two ensembles, as we show in the next section.

### E. Age-averaged protein number distribution

We would like to highlight that for discrete protein copy numbers, exact analytical number distribution in the form of series expansion has been recently obtained within the lineage ensemble [46]. Those calculations may be extended for the population ensemble, as we show below. The stationary state distributions 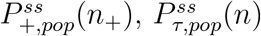 and 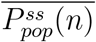 ) of protein number *n*_+_ at birth and *n* in general, follow similar equations as Eqs. III.3, III.20 and III.26, and are given below:

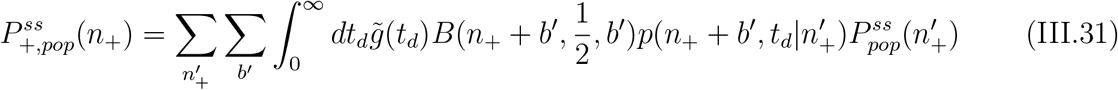

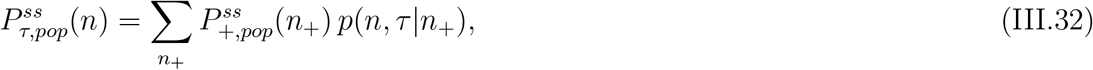

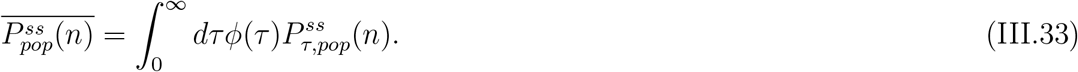

In Eq. III.31 the binomial distribution 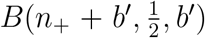 replaces the Normal distribution 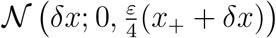 in Eq. III.3 and the integrals are replaced by discrete summations. Using the above three Eqs.III.31, III.32, and III.33, we show in Appendix. D that the generating function 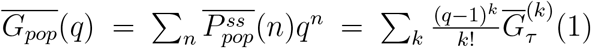 may be derived. That immediately leads to the desired explicit copy number distribution in series form:

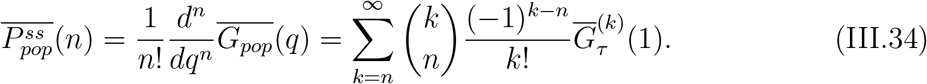

The analytical curve following Eq. III.34 is shown as a green line in Fig. 5(d) with Gillespie simulation data matching it. We also show the analytical lineage protein number distribution (brown line) alongside – the latter formula was derived in [46]. The two curves differs significantly, as the mean numbers in two ensembles are different (unlike Fig. 5(c)).

### F. Effect of correlated division times from cell-size control

The theory that we develop so far ignores inter-generational correlations for analytical tractability. But it is well known that such correlations arise due to cell-size control. Within the “adder” mechanism for example, mother-daughter generation times have negative correlation coefficient −0.25 [57, 76]. To test the robustness of our theory, we would be comparing our predictions (based on uncorrelated assumption) against the protein concentration fluctuations arising in populations with correlated division times drawn from an adder model as follows [77]. An initial cell size *s*_*b*_ grows exponentially (at rate *γ*, same as dilution rate of any protein in the cell) by adding a random amount Δ drawn from a Gamma distribution 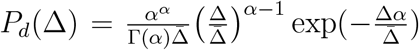 with mean 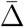, such that generation time *t*_d_ at division is given by: *s*_*b*_ + Δ = *s*_*b*_ exp(*γt*_*d*_). The initial size of the next generation 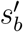 is obtained by equal partitioning of *s*_*b*_ + Δ. The evolution of protein concentration follows the kinetics already discussed above, except now following the correlated cell division times generated by the adder model. Assuming a suitable Gamma distribution for *s*_*b*_ (respecting 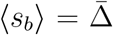 and 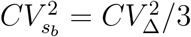) within the adder model [76], an appropriate *g*(*t*_*d*_) for the corresponding randomized division times *t*_*d*_ may be derived [46]:

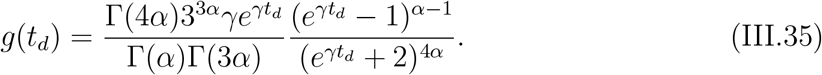

Now, if the sequence of generation times in the adder process are scrambled, then the uncorrelated times may be well represented by the distribution *g*(*t*_*d*_) in Eq. III.35. The *g*(*t*_*d*_) has two parameters *α* and *γ*. While its mean almost remains constant with *α*, it varies substantially with *γ* since ⟨*t*_*d*_⟩ ≈ (ln 2)*/γ*. On the other hand, variance of *g*(*t*_*d*_) decreases with both *α* and *γ*. Using this *g*(*t*_*d*_) in Eqs. III.9, III.16, we calculate the 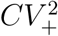 and 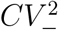 within the population ensemble, for cells which are newborn and which are about to divide. These analytical curves are plotted in Fig. 6(a-d). We observe that the variation of *CV* ^2^ is meager with *α*, in comparison with respect to *γ*.

**FIG. 6.**
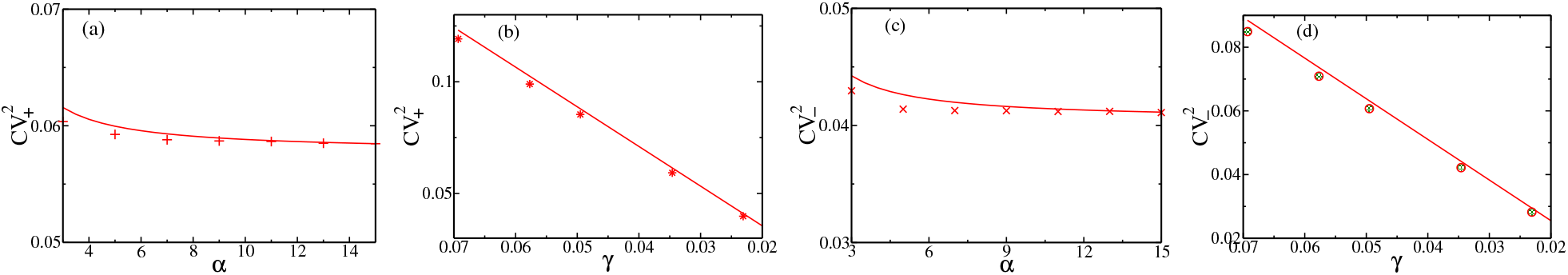
Protein concentration statistics for the adder model of cell size regulation, within a population ensemble. (a) Noise 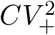 vs *α* (with *γ* = (ln 2)*/*20 min^−1^), (b) noise 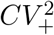 vs *γ* (with *α* = 3) for new born cells, (c) noise 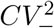 vs *α* (with *γ* = (ln 2)*/*20 min^−1^, and (d) noise 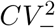 vs *γ* (with *α* = 3) for cells before division. Solid lines denote analytical results using the *g*(*t*_*d*_) of Eq. III.35, while symbols correspond to Gillespie simulations following correlated of generation time sequences from the adder model. The parameters for the protein expression are *k* = 1 min^−1^, *b* = 2 nM, *ε* = 4.0..

**FIG. 7.**
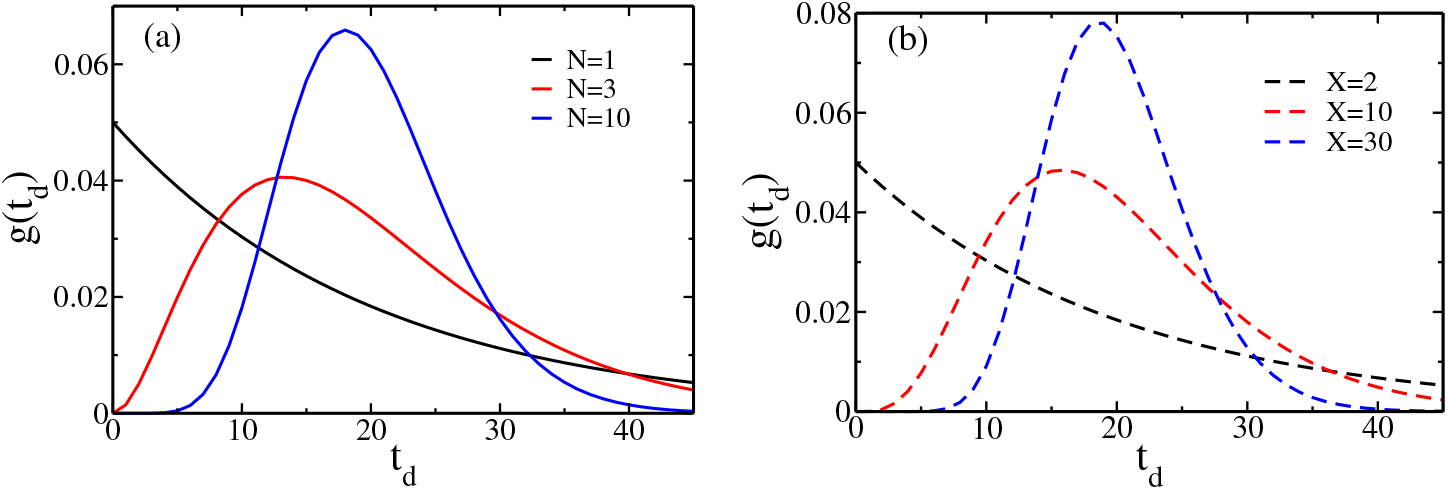
Division time distributions *g*(*t*_*d*_) are shown above. Their widths decrease (a) with the increase of the parameter *N* for the Erlang distribution, and (b) with the increase of the parameter *X* in the Beta-Exponential distributions. The ⟨*t*_*d*_⟩ = 20 min for all the distributions.

**FIG. 8.**
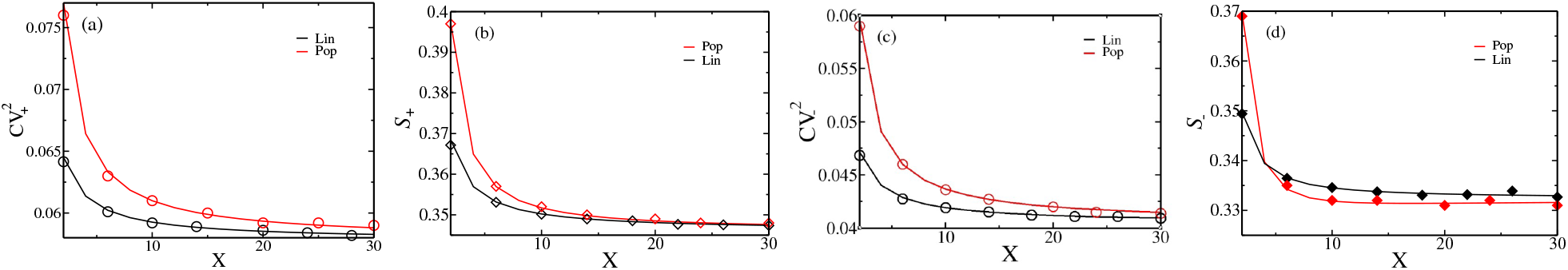
Statistics of protein concentrations (*x*_+_ and *x*_−_) for Beta-Exponentially distributed (Eq. II.3) division times. (a) Noise 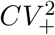 (b) skewness 𝒮_+_, (c) noise 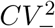 and (d) skewness 𝒮_−_ of gene products as a function of the threshold parameter *X*. Solid lines denote analytical results, while symbols correspond to simulation data. The model parameters are *k* = 1 min^−1^, *b* = 2 nM, ⟨*t*_*d*_⟩ = 20 min, *ε* = 4.0.

**FIG. 9.**
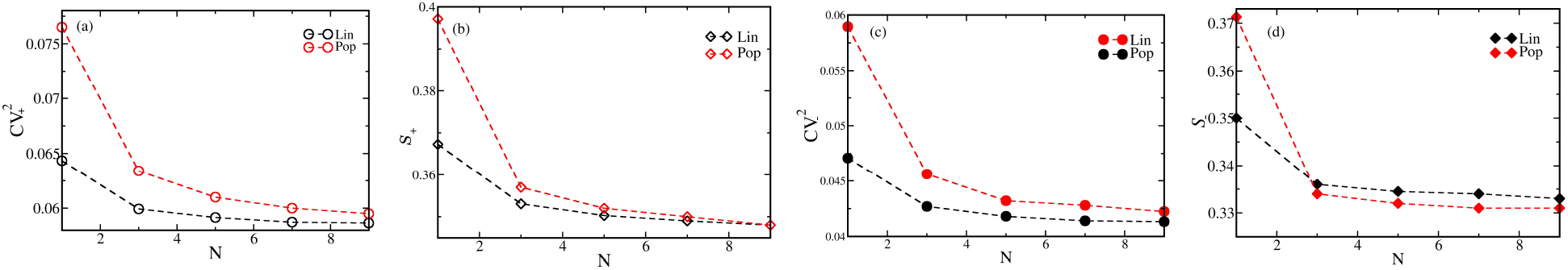
Statistics of protein concentrations (*x*_+_ and *x*_−_) for box distributed partition error is *δx* (with parameter 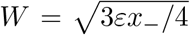, as discussed in the text in Sec.II), and with Erlang distributed division times (Eq. III.35). (a) Noise 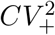 (b) skewness 𝒮_+_, (c) noise 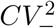 and (d) skewness 𝒮_−_ of gene products as a function of the threshold parameter *N* . The protein production model parameters are *k* = 1 min^−1^, *b* = 2 nM, ⟨*t*_*d*_⟩ = 20 min, *ε* = 4.0. All the data points are from simulations; the dashed lines join them as a visual aid.

**FIG. 10.**
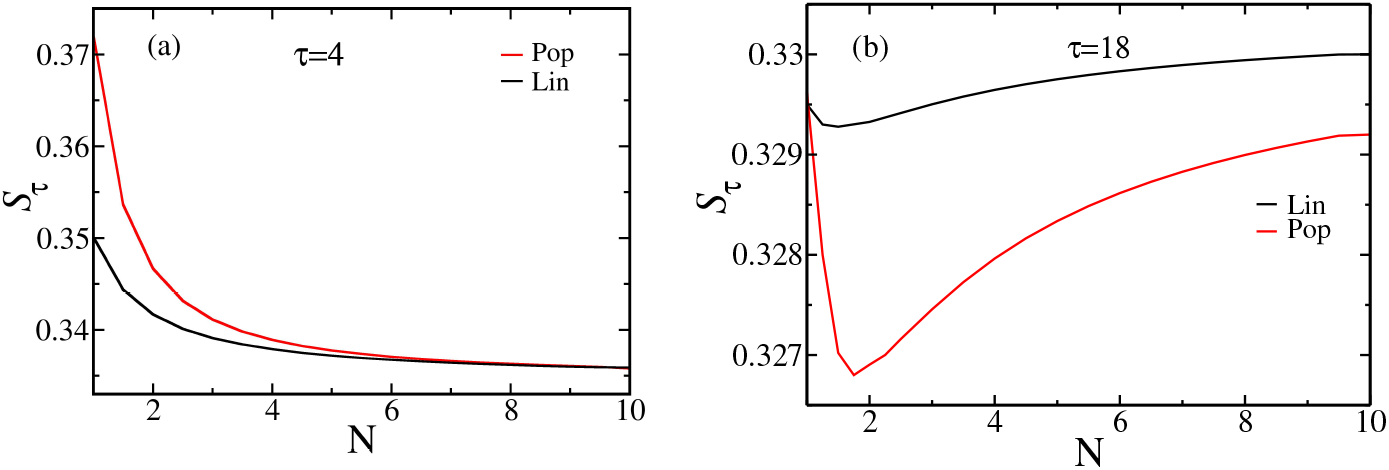
Age-specific skewness *𝒮*_*τ*_ of gene products vs *N* at cell ages (a) *τ* = 4 min and (b) *τ* = 18 min.

Using correlated generation time sequences from the adder model, we evolve the protein concentrations through Gillespie simulation, and in the cyclo-stationary state calculate 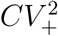 and 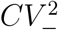. The results are shown in symbols in Fig. 6(a-d) alongside the theoretical predictions based on the assumption of uncorrelated division times. There is hardly any deviation between the correlated and uncorrelated results, except at very small *α* and *γ* (where *g*(*t*_*d*_) is quite broad). Thus inter-generational correlation with mother-daughter correlation coefficients −0.25 through cell size control by adder mechanism, is not adequately strong to cause a departure from the predictions that we have in this paper.

## IV. CONCLUSION

This work focuses on the vital topic of theory development necessary for understanding the impact of randomness in cell generation times, on the observed fluctuations in single-cell protein quantity within an ensemble, which may be of single lineages or an exponential population. Given the practical problems of drug resistance of mammalian and eukaryotic cells, these theories are hoped to be useful for the analysis of proteomic data. Recently, the issue was studied for protein copy numbers within lineage ensembles [46]. Here we generalize to accommodate the experimentally relevant feature of cell size growth and its coupling to protein concentration dilution, and derive results for the dependence of protein concentration statistics on the heterogeneity of division times.

Firstly, a striking difference is to be noted between the distributions of discrete copy numbers of proteins versus their concentrations. In the cyclo-stationary state, the *mean* concentration (*kb/γ*) does not vary at cell divisions, with change of division time distributions, nor with cell age, or between lineage or population ensembles (see curves in Fig. 2(a), 3(a), 5(c)). The full distributions of concentration are co-centered, and their differences with age and ensemble only manifest through careful study of second and higher moments and tails. In contrast, copy numbers abruptly change at every division, and as a result, distributions have clear perceptible differences between age and ensembles (see Fig. 5(d) and [46]). For this reason, from the practical applicability point of view in experiments with noisy data, full protein concentration distributions are hardly suitable to see the impact of cell division time heterogeneity. Instead, it is easier to compare the more sensitive differences of moments of concentration of different orders. In this paper, we provide moments and their exact recursion relations theoretically, to meet this relevant practical purpose. Exact formulas to all orders of moments, essentially tracks the problem mathematically formally fully, and the results are handy for data analysis. We support our analytical findings with Gillespie simulation data, and study two distinct generation time distributions *g*(*t*_*d*_), namely Erlang and Beta-exponential, as specific examples. The analytical results for Gaussian partition noise, are also compared to the simulated results for box distributed noise, and found to be very similar. We find several interesting dependencies of the moments, which we highlight below.

One of our central results is how moments depend on division time randomness at specific cell ages. Whether an experimentalist finds the concentration fluctuations in newborn or mature cells of different ages, we show that there would be a quantitatively distinct footprint of *g*(*t*_*d*_). Yet if the experimentalist samples data from all cell ages, within a lineage ensemble, that footprint vanishes at the variance level. The variance, on the other hand, remains sensitive to *g*(*t*_*d*_) within the population ensemble even if concentrations from all cell ages are considered (Fig. 5(a)). This distinction puts the population ensemble as a more suitable choice to explore division time heterogeneity. We also find that age-specific as well as age-averaged skewness in both lineage and population ensembles depend on division time randomness. Both the variance and the third cumulant decrease with a decrease in the width of the division time distributions.

Next important point is that the variance of the concentrations at any cell age (as well as averaged over age), and any *g*(*t*_*d*_) is always higher for population ensemble in comparison to the lineage ensemble (Figs. 4(a),(c), and 5(a)). Thus again population ensemble magnifies more the division time fluctuations than the lineage. On the other hand, the left-right asymmetry (through skewness) is higher for population ensemble than lineage among younger cell, and lower in its comparison in mature cells about to divide (Figs. 4(b),(d) and Fig. 10). This interesting feature gets washed away for age-averaged skewness (Fig. 5(b)).

While inter-generational correlations indeed are present in cell generation times, the degree of mother-daughter correlation co-efficient of −0.25 is insufficient to have any significant impact on concentration fluctuations, as compared to independent generation times. We have explored this with varying cell dilution rate *γ* and shape parameter *α* (see Fig. 6). This is similar to the earlier findings on number fluctuations [46].

The initiative which started recently to find protein number and distributions for arbitrary random cell generation times, got extended through this work to include effects of cell growth. The aspect of gene dosage with associated DNA duplication, remains to be incorporated further in the picture, to make it more complete in future studies.

## ethics

This work did not require ethical approval from a human subject or animal welfare committee.

## contribute

S.Y.A.: conceptualization, methodology, software, data curation, formal analysis, investigation, writing—original draft, writing—review and editing; A.P: conceptualization, supervision, writing—review and editing; A.S: conceptualization, resources, supervision, writing—review and editing; D.D: conceptualization, methodology, data curation, formal analysis, investigation, writing—original draft, writing—review and editing, and supervision. All authors gave final approval for publication and agreed to be held accountable for the work performed therein.

## competing

We declare we have no competing interests.

## acknowledgments

DD acknowledges the visitor program of MPI-PKS Dresden (where a part of this work was done in summer 2025). SYA acknowledges IIT Bombay for financial support through the institute’s post-doctoral fellowship.

## disclaimer

We have not used AI-assisted technologies in creating this article.

## Appendix A: Appendix A: Division time Distributions

Here we show the shapes of the division time distributions that we used in the main text.

## Appendix B: Appendix B: Concentration distributions and general moments in newborn cells

The probability distributions of protein concentrations in newborn cells in the *i*th and (*i* − 1)th cell cycle are related as follows

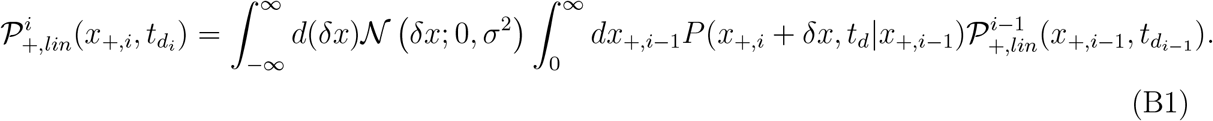

Multiplying the above equation with 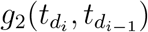 and assuming uncorrelatedness of the division times 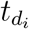 and 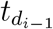, and defining 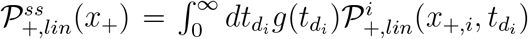 for *i* ≫ 1 we obtain Eq. III.1 (same as Eq. B2):

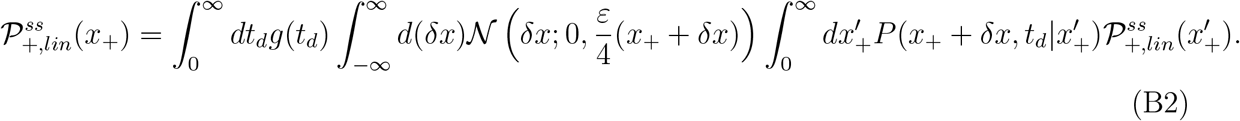

The forward master equation for the probability distribution of protein concentration 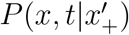 with 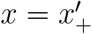 at *t* = 0, is given in Eq. II.1. Taking Laplace transform 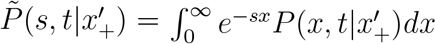, and using the method of Lagrange characteristics, one obtains

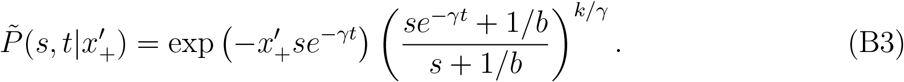

which may be used in the following calculation.

Taking the Laplace transform of 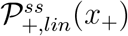 in Eq. B2, using Eq. B3, and noting 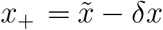 (with 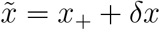) and 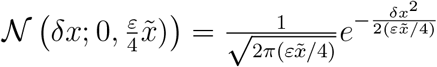 we get Eq. III.2 (same as Eq.B4):

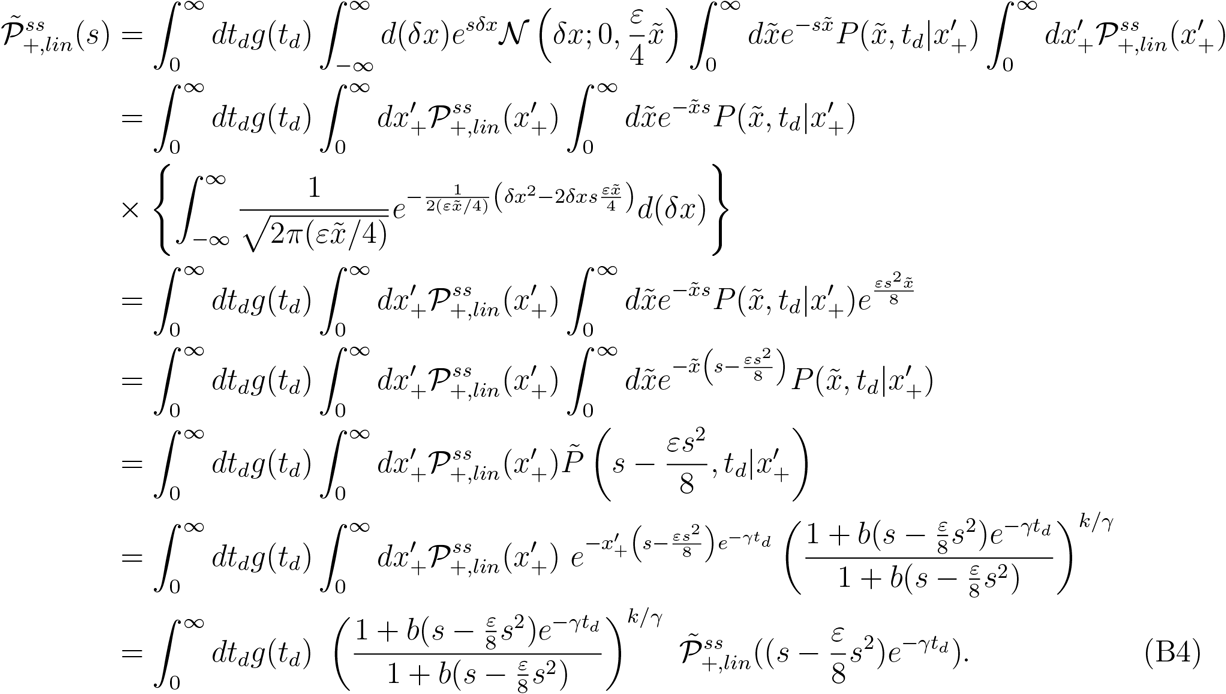

The approximation 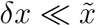 is assumed valid for the above calculation to be justified physically, preventing in practice the Gaussian tail from extending into the non-physical negative concentration domain.

Starting from a similar equation for 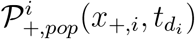 as Eq. B1 with the extra factor 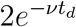 on the right hand side, we obtain Eq. III.3 in the cyclo-stationary state for the population ensemble. Taking the Laplace transform of 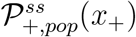 in Eq. III.3 and using Eq. B3 just as above, we get Eq. III.5.

The Eq. III.2 (or Eq.B4) may be rewritten as below, using the inverse mapping *u*(*s*^*′*^) of 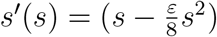:

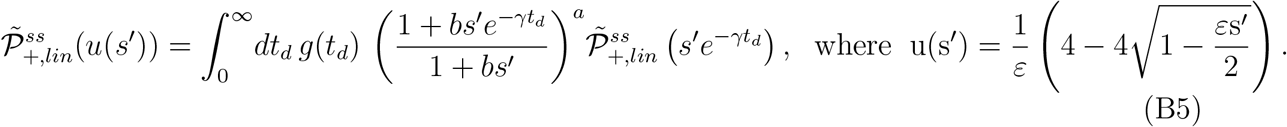

Starting from Eq. B5 we may Taylor expand the left hand side about *s*^*′*^ = 0 as 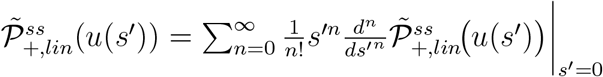, where the *n*-th derivative is given by Faà di Bruno’s formula:

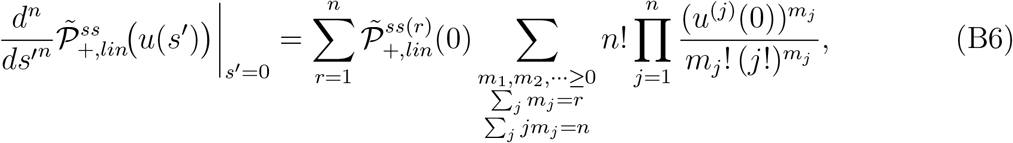

Where 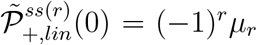, from Eq. III.6. Similarly, expanding the *s*^*′*^ dependent function in the integrand on the right-hand side of Eq. B5 and using Eq. III.6, we finally obtain the Eq. III.7 by equating the coefficients of *s*^*′n*^ on both sides.

A separate but similar treatment can be done for the population ensemble, as mentioned in the main text.

## Appendix C: Appendix C: Concentration distributions of cell about to divide

The pre-division distribution 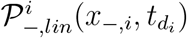 at the end of (*i* − 1)the cycle is related to newborn distribution 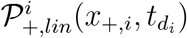 in the *i*th cycle as follows

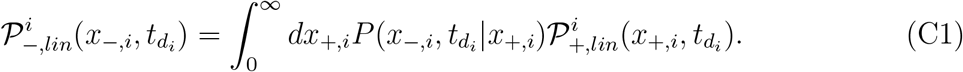

Defining 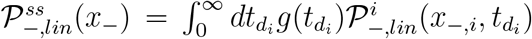 for *i* ≫ 1 we obtain Eq. III.11 for the stationary concentration distribution just before cell division within lineage ensemble. The corresponding equation within population ensemble appears in Eq. III.12. The Laplace transform of the Eq. III.11 is

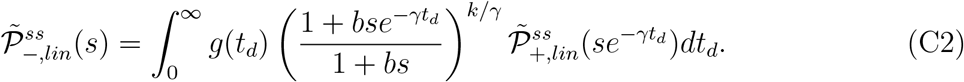

Comparing the above equation with Eq. B5 we get

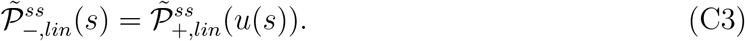

which is Eq. III.13. The same argument applies for population ensemble.

## Appendix D: Appendix D: Cell age-specific moments

Taking the Laplace transforms of Eq. III.19 and III.20 and substituting in those, the form of 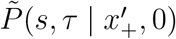 from Eq. (B3), we get

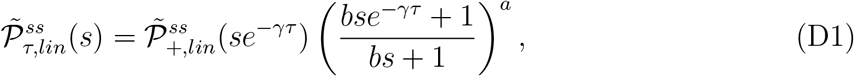

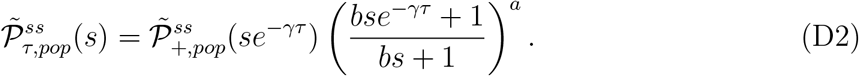

Expanding both sides of above two equations as a series in *s*, and equating the coefficients of *s*^*n*^ we get any desired moment of 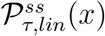 and 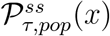:

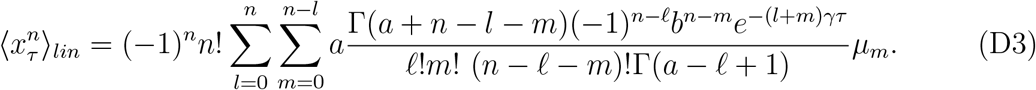

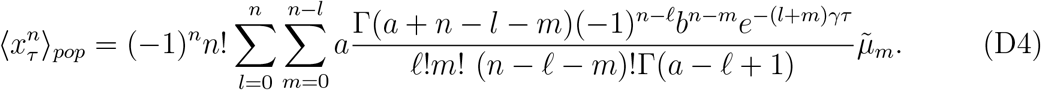

Note for the population ensemble *µ*_*m*_ is replaced by 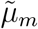.

In Fig. 4(b) we have shown that at cell age *τ* = 10 min, 𝒮_*τ*_ vs *N* shows a crossing of the population and the lineage curves. At the same time we saw that in Fig. 2(d) for the newborn cells there is no such crossing of 𝒮_+_, while for cells prior to division in Fig. 3(d) 𝒮_−_ shows a crossing. We studied the range of *τ* for which this crossing is present and found that to be *τ* ∈ (4, 18). We show here (Fig. 10) that at *τ* = 4 min the 𝒮_*τ*_ curve for population is entirely above lineage, while for *τ* = 18 min their positions are reversed.

## Appendix E: Appendix E: Cell age-averaged moments

From Eqs. III.25 and III.26, the Laplace transforms 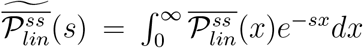 and 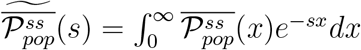 are given by:

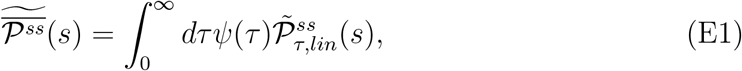

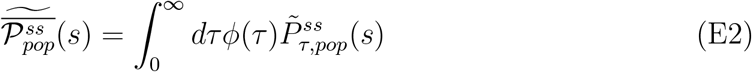

Substituting Eqs. D1 and D2 in Eqs. E1 and E2 respectively, series expanding in *s*, and equating coefficients of *s*^*n*^ from both sides, we obtain the exact *n*th moments of the age-averaged distributions in the two ensembles:

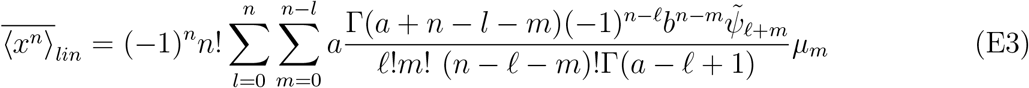

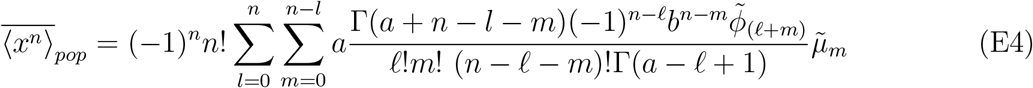

## Appendix F: Appendix F: Protein copy number distribution in population ensemble

The stationary state distribution 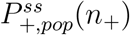, of protein number *n*_+_ at birth within a population ensemble follows Eq. III.31. Its generating function is

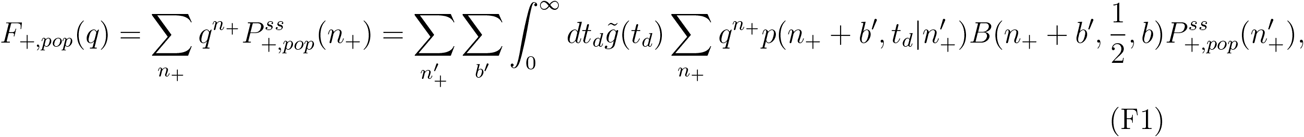

where 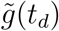 is given by Eq. III.4. After few steps of algebra, it reduces to

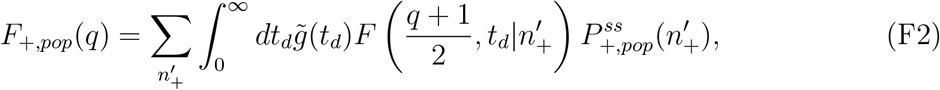

where the generating function 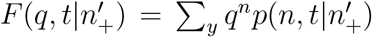 for bursty protein expression with degradation rate *γ*_*p*_ is given by [25, 29]:

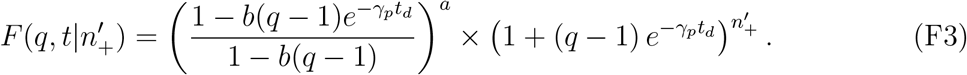

Substituting the above in Eq. F2, and expressing the generating function as a series 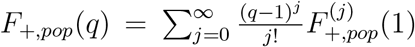 about *q* = 1 on both sides, we have the recursion relation for the coefficients 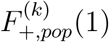. in Eq. F4:

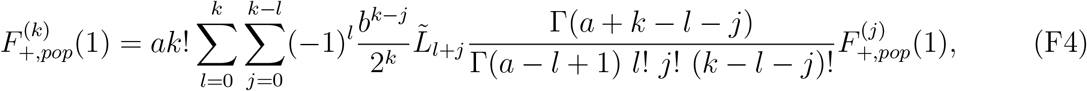

with 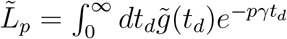

Next starting from the Eq. III.32 for the age-dependent distribution 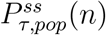 and defining the generating function 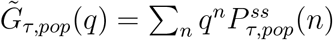 we get

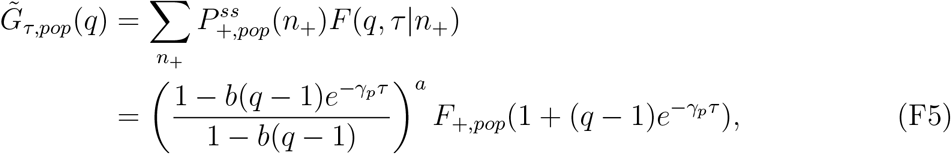

where we have used Eq. F3 and the definition of *F*_+,*pop*_(*q*) from Eq. F1. Further defining 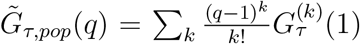, expanding the right hand side of Eq. F5 in a power series of (*q* − 1), and comparing the coefficients on both side we get

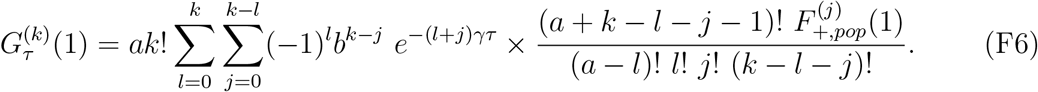

Finally we note that the age-averaged protein concentration distribution 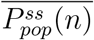 is related to age-specific distribution 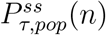 through the Eq. III.33. Using the generating functions 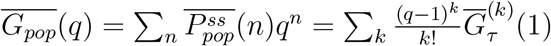 and 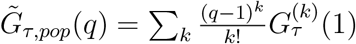, we obtain

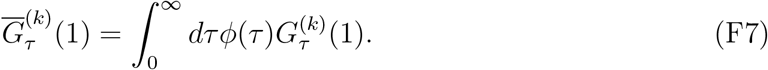

Using Eq. F6 in Eq. F7 we get 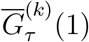 in Eq. F8:

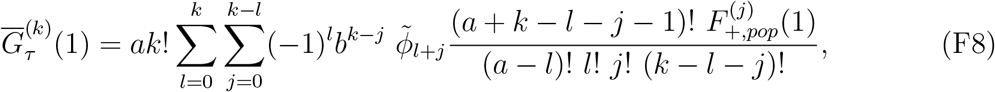

where 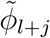 is given in Eq. E4. This further leads to the explicit age-averaged protein number distribution 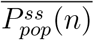 in the population ensemble given by Eq. III.34.

